# Moisture adsorption isotherms and quality of seeds stored in conventional packaging materials and hermetic Super Bag

**DOI:** 10.1101/463026

**Authors:** Muhammad Amir Bakhtavar, Irfan Afzal, Shahzad Maqsood Ahmed Basra

**Author notes:** **Corresponding author**, /2949. These authors contributed equally to this work (Muhammad Amir Bakhtavar, Irfan Afzal).

## Abstract

Seed moisture content (SMC) is an important attribute to seed quality. Maintaining seed dryness throughout supply chain (The Dry Chain) prevents seed germination and quality losses. Ambient relative humidity (RH) and temperature affect seed moisture and thereof seed moisture isotherm. Present study was conducted to compare the moisture adsorption isotherms of wheat, maize, cotton and quinoa seeds packed in hermetic Super Bag and traditional packaging materials including paper, polypropylene (PP), jute and cloth bags. Seeds were incubated at 60, 70, 80 and 90% static RH. Nearly straight line moisture isotherms for all crop seeds were obtained in Super Bag. Seed moisture contents increased in traditional packaging materials with increasing RH. At higher level of RH, moisture contents increased slightly (1-2%) in Super Bag, whereas this increase was much higher in traditional packaging materials (≈9% higher than original SMC at 90% RH). In second study, seeds were dried to 8 and 14% initial seed moisture contents using zeolite drying beads and were stored in hermetic and traditional bags for a period of 18 months. For all crop seeds, germination was severely affected in all packaging materials both at 8 and 14% initial SMC except storage in Super Bag at 8% SMC. Wheat seed stored in Super Bag at 8% SMC almost maintained initial germination while germination of cotton, maize and quinoa seeds declined 7%, 14% and 30% respectively in Super Bag at 8% SMC. Seed storage in Super Bag can help to prevent the significant increase in seed moisture at higher RH as is evident from moisture isotherm study, thus helps to preserve quality of maize, wheat, cotton and quinoa seeds by maintaining The Dry Chain throughout the storage period.

## Introduction

Seed moisture is a critical factor influencing seed quality, as seed shelf life is highly dependent upon its moisture contents. Knowledge of best seed moisture content for seed storage both increases shelf life and reduces contamination by storage fungi [1]. Knowing seed moisture even in the field of seed testing is essential to avoid the imbibitional injury during germination process [2] and it is essential parameter in standardizing results of vigor test such as conductivity test and accelerated aging test [3]. Bewley et al. [4] emphasized that seed quality is at greatest risk at high moisture content during storage. Rate of seed reparation increased at high seed moisture contents and heat generated by the respiring seeds is enough to kill the seeds [5]. Insect infestation is minimal below 8-9% moisture content. A non-dormant seed may germinate above 30% seed moisture content. Fungi cannot grow below 13% seed moisture contents in starchy seeds and 7-8% in oily seeds which are in equilibrium with ambient RH values nearly equals to 68% [1, 4]. Storage of maize at high moisture contents (15%) resulted into low germination percentage, dry matter losses (up to 35%) along with fungal growth [6].

Relationship between prevailing RH and SMC at a given temperature is known as seed moisture isotherm [7]. Seed moisture adsorption isotherms are an important tool to predict the potential changes occurring in the biological material’s stability [8]. Seed is a hygroscopic entity and absorbs moisture from the surrounding, so any change in temperature and RH of the surrounding, affects its moisture isotherm, moisture contents and ultimately its quality [9]. Nature of the packaging materials has great influence over the seed moisture contents. Different packaging materials have different water vapor transmission rates [10] thus will have difference in the moisture contents of seeds contained inside them. In most of south Asian countries including Pakistan, seeds are stored in conventional porous bags and earthen bins [11]. Large pore size of jute, cloth and polypropylene bags provide free access to the water vapors that were readily absorbed by the seeds and ultimately elevated seed moisture contents.

Maintaining seed dryness by implementing The Dry Chain reduce storage losses of stored commodities [1] and offers an opportunity for safe storage of seeds without significant loss of germination. Main idea behind dry Chain concept is initial drying of seeds/grains to low moisture contents and then maintenance of that dryness throughout supply chain. There is no study available on seed moisture isotherm variations within conventional porous bags with changing RH and its ultimate effect on seed quality. The present study was conducted to compare seed moisture adsorption isotherm of different crop seeds at various levels of RH when stored in conventional porous packaging materials and hermetic Super Bags. Moreover, germination of wheat, maize, cotton and quinoa seeds stored in these packaging materials at 8 and 14% initial seed moisture contents, after 18 months storage have been compared to check if maintaining seed dryness (The Dry Chain) could be a strategy to preserve seed quality.

## Materials and methods

### Seed moisture isotherm study

Experiment was performed in Seed Physiology Lab, Department of Agronomy, University of Agriculture Faisalabad, Pakistan. Hermetic plastic Super Bags were provided by the GrainPro Inc. USA and conventional porous packaging materials (paper, PP (Poly propylene), jute and cloth bags) used in this study were purchased from local grain market. All the bags were cut into smaller size having capacity of 150 g seeds and edges were sealed with scotch tape. Hermetic Super Bags were cut in a way that only bottom portion of the bag was used so as to maintain its hermetic nature. After determination of initial moisture contents on fresh weight basis, wheat (9%), maize (9.2%), quinoa (7.5%) and cotton (7.65%) seeds were packed in different packaging materials. After packaging seeds were incubated at four static RH levels (60, 70, 80 and 90%) maintained in airtight plastic boxes keeping a constant temperature of 25°C.

### Maintaining static RH levels

Saturated salt solutions were used to maintain static RH levels in airtight plastic boxes. Saturated solution of FeCl_2_.4H_2_O gave 60% RH and NaCl+NaNO_3_ maintained ≈70% at 25°C. Similarly saturated solution of NaH_2_PO_4_ maintained 80% RH while BaCl_2_.2H_2_O gave 90% RH at 25°C [12].

### Determination of seed moisture contents

Seed moisture contents of each crop were recorded after 15 days of incubation at each of the four RH levels (60, 70, 80 and 90%). Four replications of ≈ 5 g seeds were dried in an oven at 103°C for 17 hours [13] and dry weight of each replicate was recorded to calculate the seed moisture contents on fresh weight basis.

### Development of moisture isotherms

Moisture isotherms were developed by plotting the data of seed moisture contents against different RH levels (60, 70, 80 and 90%) in the form of a line graphs in Microsoft Excel.

### Seed storage study

Seed storage experiment was performed in Seed Physiology Lab, Department of Agronomy, University of Agriculture Faisalabad, Pakistan. Seeds of wheat cv. Galaxy 2013 and cotton cv. Lalazar were obtained from Ayub Agriculture Research Institute, Faisalabad, Pakistan. Wheat seed’s initial germination was 99.8% whereas initial germination of cottonseed was 75%. Hybrid maize (30Y87) seeds were provided by Pioneer Pakistan Seeds Ltd. Sahiwal, having 99% initial germination. Quinoa (Cv. UAFQ7) seeds were procured form Crop Physiology Research Area, Department of Agronomy having initial 80.5% germination.

### Seed drying

Maize and wheat seeds (5 kg per replicate) were dried to 8% seed moisture contents by mixing with zeolite seed drying beads in an airtight container. Drying beads (Rhino Research Group Thailand) are made up of aluminium silicate ceramics (Zeolite clay) material with very small, uniform pores where water molecules can be adsorbed and are used for quick drying of seeds. Amount of beads required to dry seeds was calculated using a bead calculator [1].

### Equilibrating seed moisture contents

Maize seeds were incubated over saturated salt solution of FeCl_2_ for 20 days to maintain 14% equilibrium seed moisture contents at 25°C. To maintain 14% moisture contents, wheat seeds were incubated over saturated salt solution of NaNO_2_ at 25°C [12] for 20 days. Cotton seeds were incubated at saturated salt solution of Mg(NO_3_)_2_ and BaCl_2_ to get 8% and 14% equilibrium seed moisture contents respectively. Similarly, quinoa seeds were incubated over saturated salt solution of CaCl_2_ and CuBr_2_ at 25°C for 20 days in airtight plastic box to get 8% and 14% seed moisture levels respectively. Seeds of each crop equilibrated to 8% and 14% seed moisture levels were packed in Super Bag, paper, PP, jute and cloth bags and were stored for 18 months in Seed Physiology Lab at ambient conditions.

### Storage environment’s temperature and relative humidity data

Data of temperature and RH for whole storage duration (July 2015 to July 2017) were recorded with the help of Data logger (Centor Thai, Thailand) that was installed in the store house. During 2015, maximum RH was recorded in the month of November while minimum RH was recorded during October (Table 1). Maximum temperature was recorded during the month of July whereas minimum temperature was recorded in December. In 2016, maximum relative humidity was recorded in the months of January and February while minimum relative humidity was recorded during May. Maximum temperature was recorded during the month of May whereas minimum temperature was recorded in January. During 2017, maximum RH was recorded in the month of March while minimum RH was recorded during May. Maximum temperature was recorded during the month of June whereas minimum temperature was recorded in January (Table 1).

**Table 1.**
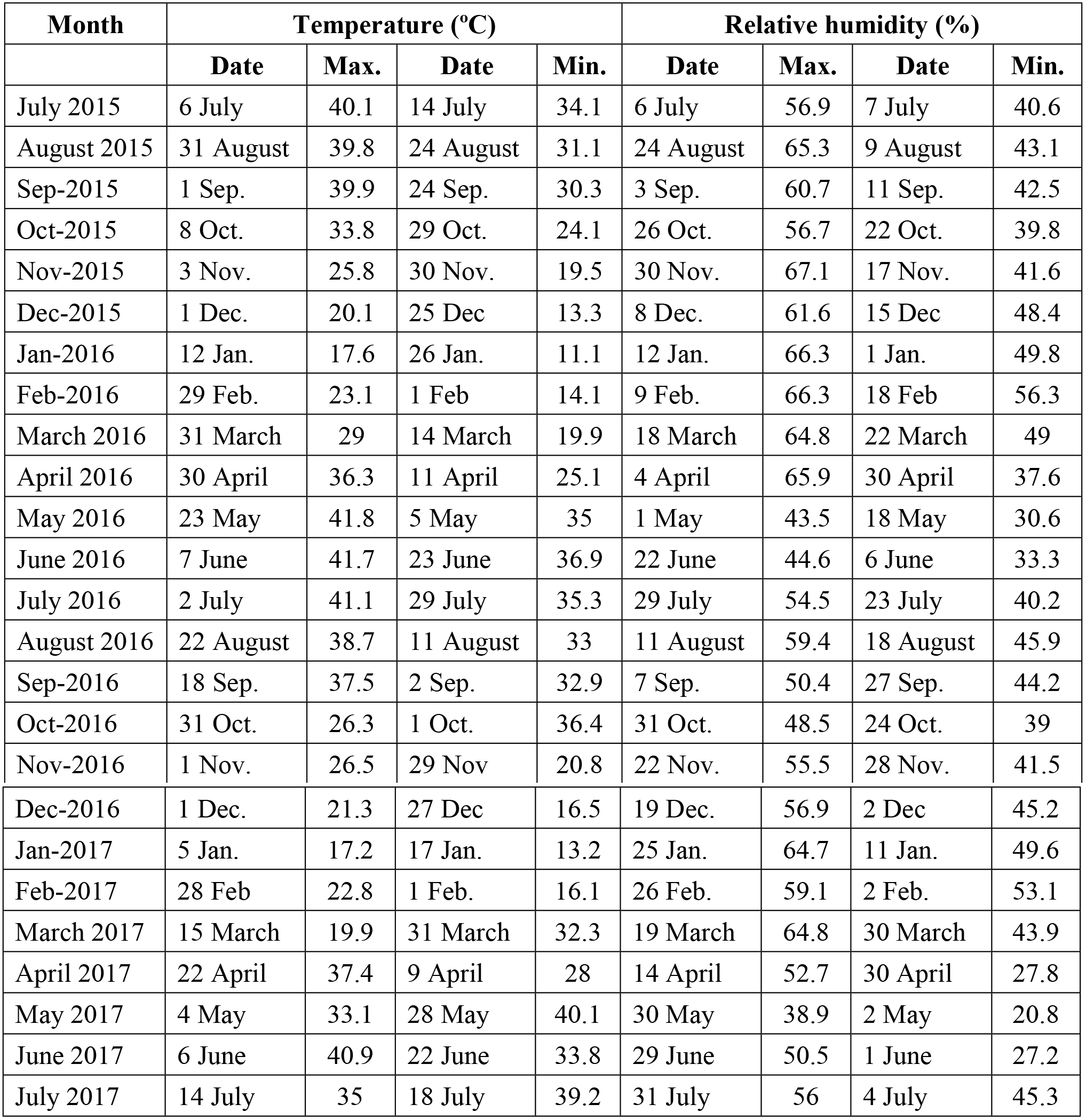
Monthly maximum and minimum values of temperature and relative humidity recorded with data logger installed in seed store house.

### Seed germination testing

Seed sampling was done after 18 months of storage. Seed germination was tested by placing 100 seeds in sterilized and well moist blotting paper and making four similar replications to test total 400 seeds from randomly drawn seed samples in a germinator (SANYO Japan, MIR-254) at 25°C [13]. Seedlings were evaluated according to ISTA Handbook for seedling evaluation [14] and percentage of normal seedlings was reported as final germination.

### Statistical analysis

Completely Randomized Design with factorial arrangements was employed for data analysis using statistical software Statistix 8.1. Seed moisture contents and packaging material were treated as factors and each experimental unit was replicated three times. Treatment means were compared using Tukey’s Test at P ≤ 0.05.

## Results

### Moisture Isotherm study

Cottonseed having highest oil contents on percent weight basis had lowest seed moisture contents at all levels of RH (Fig 1). At 20% RH wheat seeds have highest moisture contents (6.1%) whereas cottonseeds have lowest moisture contents (4.25%). Similarly, cottonseeds have lowest moisture at 90% RH and wheat seeds have highest moisture contents at this humidity level. Maize and quinoa seeds have moisture contents higher than cotton seeds with moisture contents more close to the wheat seeds at all levels of RH and slight overlapping at some points (Fig 1).

**Fig 1.**
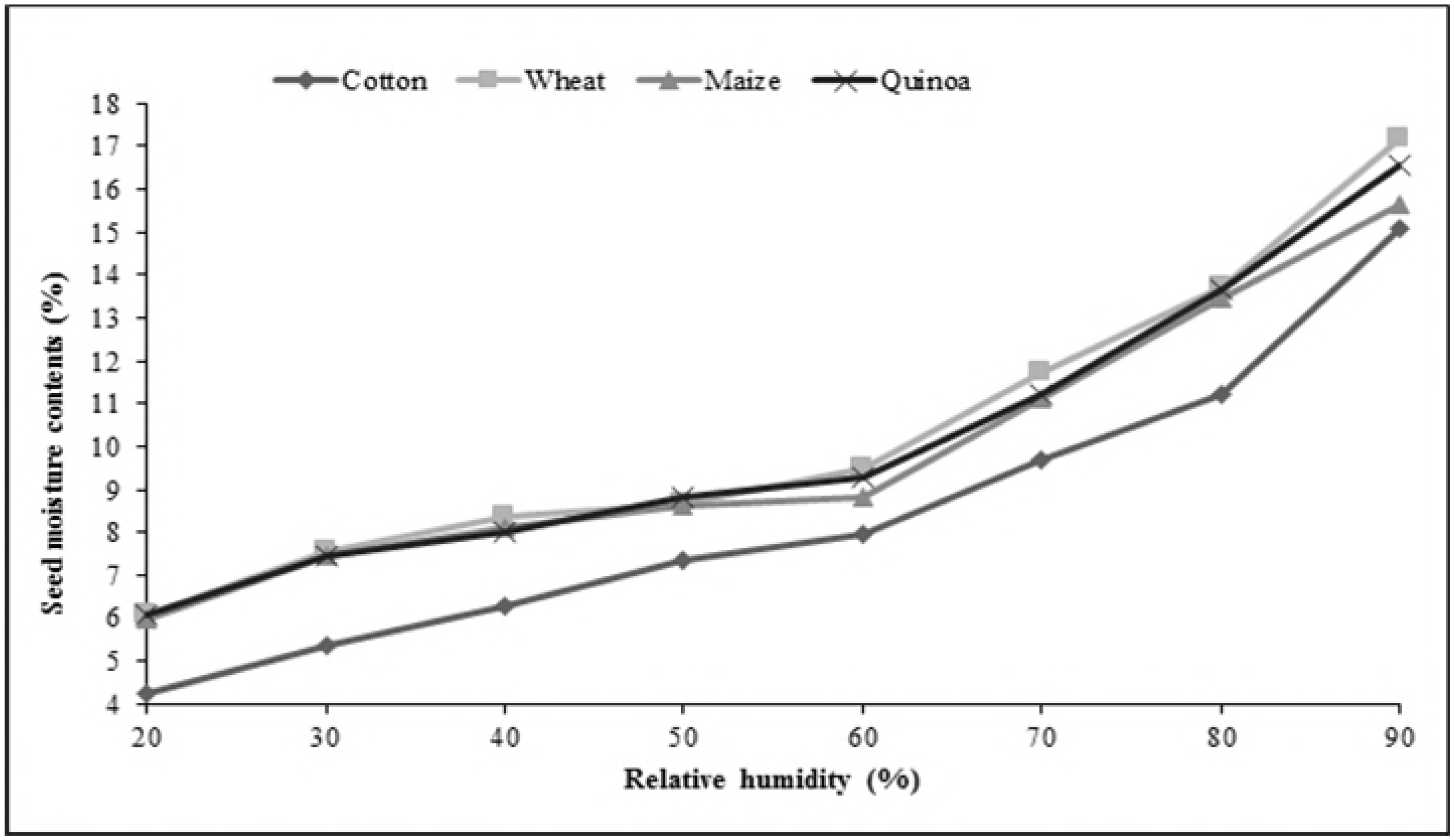
SeEd moisture adsorption isotherms of cermls, psudocereals and oilseeds at 25°C. Within different packaging materials, maize seed had moisture contents in range of 9.99% (Super bag) to 10.56% (Cloth bag) at 60% RH. There was no significant difference in moisture contents of maize seed placed in conventional packaging materials at 60% RH (Fig 2). Similarly, moisture contents were lower in the seeds when placed in Super bag and paper bag at 70% RH while rest all bags have high seed moisture. At 80% RH, Super Bag maintained minimum seed moisture contents while seed in cloth and PP had maximum seed moisture contents. At 90% RH, although seed moisture contents increased from 10% (at 80% RH) to 10.89% in Super Bag but this increase was far less than that was observed for seed moisture in other packaging materials. Seed in cloth bag had maximum seed moisture contents (15.86%). Overall, moisture isotherms indicate that Super Bag maintained seed moisture at all levels of RH while seed moisture contents varied linearly with prevailing RH in all bags especially in cloth, PP and jute bag (Fig 2).

**Fig 2.**
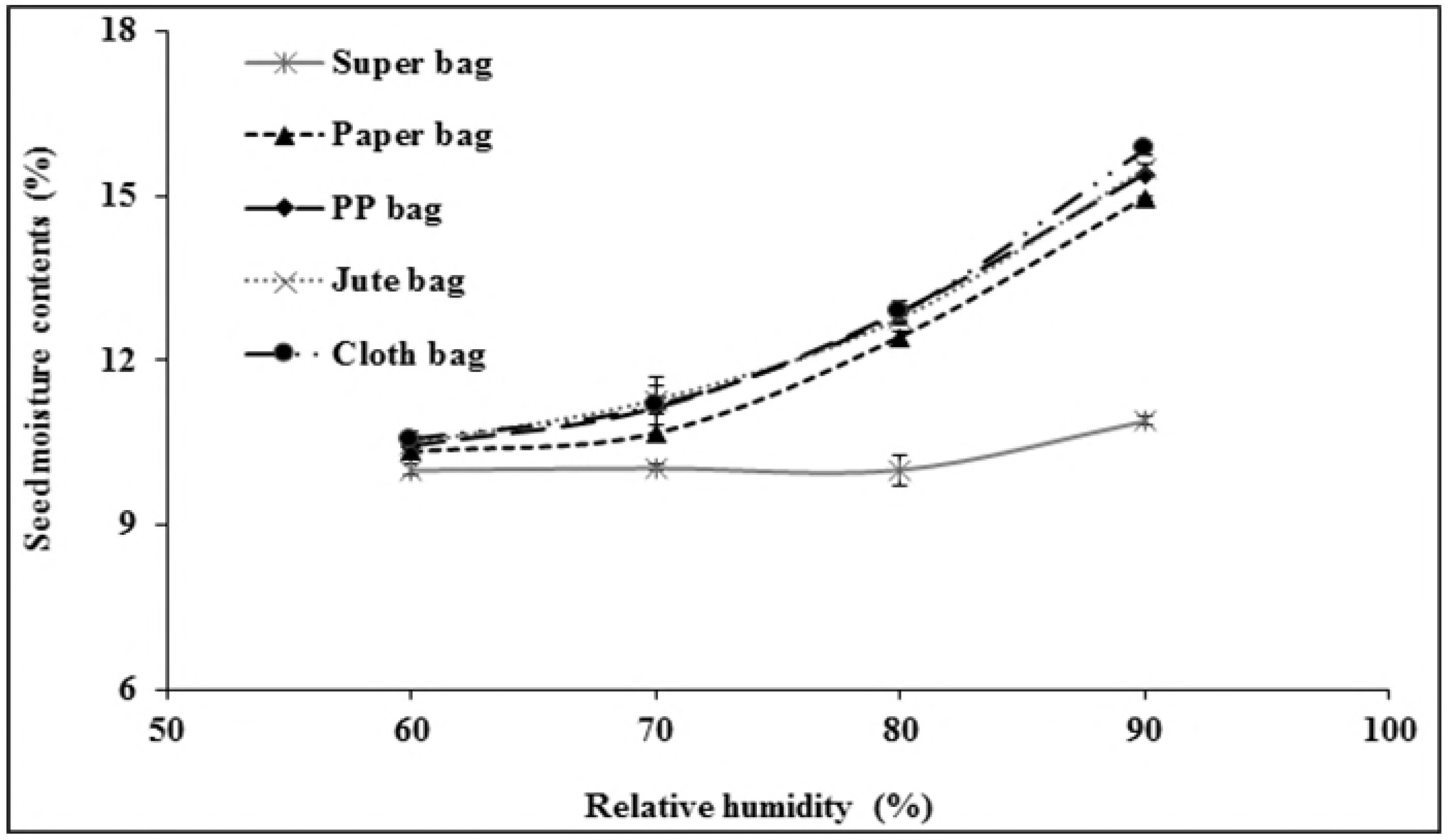
Seed moisture adsorption isotherms of maize seeds in different packaging materials. at 25°C. There was no significant difference in moisture contents of wheat in all packaging materials at 60% RH (Fig 3). At 70% RH, Super Bag and paper bag had seeds with lowest moisture contents while seed in cloth and jute bag had higher moisture. Moisture contents of wheat seeds were strikingly different in various packaging materials at 80% RH. Super Bag had seed with minimum moisture contents followed by seeds moisture in paper bag. Rest all bags had higher seed moisture contents. At 90% RH, there was abrupt increase in moisture contents of wheat seed in Super Bag as well. Seed moisture contents increased from 9.69% (at 80% RH) to 10.32% in Super Bag. Seed in cloth bag had maximum seed moisture (17.1%).

### Seed storage study

Seed germination was highly linked with the initial seed moisture contents and storage conditions. Moreover, type of packaging materials also significantly affected seed germination of all crops under study. Highest germination (85%) of maize seed was recorded when it was stored in Super Bag at 8% initial seed moisture contents (SMC). Even in Super Bag, maize seed lost its germination completely when it was stored at 14% initial SMC. Likewise seed germination after 18 months of storage was drastically reduced in all conventional packaging materials (Paper, PP, jute and cloth bag) irrespective of initial seed moisture contents (Table 2). Wheat seed germination was fairly good in all packaging materials with maximum (99%) germination in Super Bag after 18 months storage at 8% initial seed moisture contents. Wheat seeds stored in all other conventional packaging materials at 8% initial SMC also had statistically similar germination percentage. Wheat seed stored in Super bag at 14% initial SMC had lowest (18%) germination, however, germination declined only slightly in conventional bags when stored at 14% initial SMC (Table 2).

**Table 2.** Final germination of crop seeds after 18 months of storage in different packaging materials at various levels of initial seed moisture contents.

| Packaging Materials | Initial Germination (%) | Final Germination at 8% SMC | Final Germination at 14% SMC | Initial Germination (%) | Final Germination at 8% SMC | Final Germination at 14% SMC |
| --- | --- | --- | --- | --- | --- | --- |
|  | Maize |  |  | Wheat |  |  |
| Super Bag | 99 | 85±2.89a<sup>z</sup> | 0±0 b | 99.8 | 99±0.50 a | 18±1.67 b |
| Paper bag | 99 | 8±1.67 b | 0±0 b | 99.8 | 95±5.01 a | 88±4.41 a |
| PP bag | 99 | 3±1.67 b | 2±2.89 b | 99.8 | 88±2.89 a | 82±1.67 a |
| Jute Bag | 99 | 3±2.89 b | 2±1.67 b | 99.8 | 85±11.68 a | 78±7.65 a |
| Cloth Bag | 99 | 8±4.42 b | 2±1.67 b | 99.8 | 87±3.34 a | 83±8.83 a |
| HSD at P ≤ 0.05 | 11 |  |  | 31 |  |  |
|  | Quinoa |  |  | Cotton |  |  |
| Super Bag | 80.5 | 50±2.89 a | 0±0 d | 75 | 69±0.58 a | 0±0 d |
| Paper bag | 80.5 | 30±1.73 b | 16±1.67 c | 75 | 62±0.88ab | 7±2.41 d |
| PP bag | 80.5 | 23±4.41bc | 17±1.53 c | 75 | 58± 0.67b | 28±3.47 c |
| Jute Bag | 80.5 | 21±3.34bc | 18±3.34 bc | 75 | 59±1 b | 36±1.16 b |
| Cloth Bag | 80.5 | 23±1.67bc | 20±2.89bc | 75 | 60±0.66ab | 29±3.53 c |
| HSD at P ≤ 0.05 | 12 |  |  | 9 |  |  |

Means within same column not sharing same letters are significantly different at $p \leq 0.05$ . PP; Polypropylene, SMC; Seed moisture contents.
<sup>z</sup>Mean separation within columns by Tukey's HSD at $p \leq 0.05$ .

Cotton seed stored in Super Bag at 8% initial SMC gave maximum germination (68%) followed by germination (62%) of seed stored in paper bag after 18 months. Minimum seed germination was recorded when cotton seed was stored in Super Bag at 14% initial seed moisture contents. Seed storage in conventional porous packaging materials at 8% initial seed moisture contents, resulted into ≈ 15% losses in germination while these losses were higher (40-60%) when stored at 14% initial seed moisture contents (Table 2). Quinoa seed germination was drastically affected in all packaging materials at both 8 and 14% seed moisture contents. After 18 months storage, highest germination (50%) was recorded for the seeds stored in Super Bag at 8% initial SMC. Seed storage in conventional packaging materials at 8% initial SMC, resulted into 50-60% losses in germination as compared to germination losses (30%) in Super Bag at 8% initial SMC. Quinoa seed completely lost its germination when stored in Super Bag at 14% initial SMC.

## Discussion

Due to its hygroscopic nature seed absorbs and desorbs moisture according to ambient relative humidity and attain equilibrium [9]. Seed moisture contents vary continuously according to RH and temperature of the environment. Extremely low water potential within the dry seeds creates greater affinity for the water molecule to readily absorb by the seeds. Seed moisture isotherms describe equilibrium relationship between the seed moisture contents and equilibrium RH at a given temperature [15]. If storage environment is a closed container, the seeds tend to equilibrate itself with that microclimate and RH of that closed container is known as equilibrium relative 14 humidity. Seed moisture contents varied in different packaging materials due to varying water vapor transmission rates [10]. For all crop seeds, Super Bag had lower moisture contents at all levels of RH (Figs 2-5). This low moisture content is due to the hermetic nature of the Super Bag that created hindrance to the incoming water vapors even at higher RH. Super Bag is made up of multilayers of polyethylene having a less permeable barrier layer to prevent exchange of moisture and air and has very low vapour transmission rate i.e. <10 gm-2day-1 [16]. Seed moisture increased slightly in Super Bag at higher RH levels (80 and 90%).

Moisture contents in all conventional bags varied according to the ambient RH and continuously increased with increasing RH. Most of the crop seeds had highest moisture in cloth bag. Free exchange of moisture between seed and environment resulted into high moisture of the seeds at higher RH (Figs 2-5). Likewise, seed moisture contents were also higher when seeds were packed in jute and cloth bags. Large pore size of the jute, cloth and polypropylene bags provided free access to the water vapors which were readily absorbed by the seeds and ultimately elevated seed moisture contents.

Seed moisture isotherm also varies with seed oil contents [17]. Results of current study indicated that at a given level of RH there was a difference in seed moisture contents of different crops in same bag. For example at 90% RH, moisture contents of wheat seeds were 16.35, 17.10, and 16.83 whereas moisture contents of maize seeds were 15.86, 15.50 and 15.39 % in cloth, jute and PP bag respectively (Figs 2 and 3). This difference in seed moisture contents for both crop seeds is due to difference in seed composition as wheat seed has ≈2.3% oil contents and 12% protein contents [18] whereas maize seed has 5% oil contents and 9% protein contents [19]. Wheat seed has more protein contents and protein has greater affinity for water molecule due to presence of positive and negative charges that provide site for hydrogen bonding [17] thus had higher moisture contents. Moreover, maize seeds have more oil contents as compared to wheat seeds and lipids have less affinity for the water molecule so low oil contents and higher protein contents were responsible for the elevated moisture contents in wheat seeds in this study (Figs 2 and 3).

**Fig 3.**
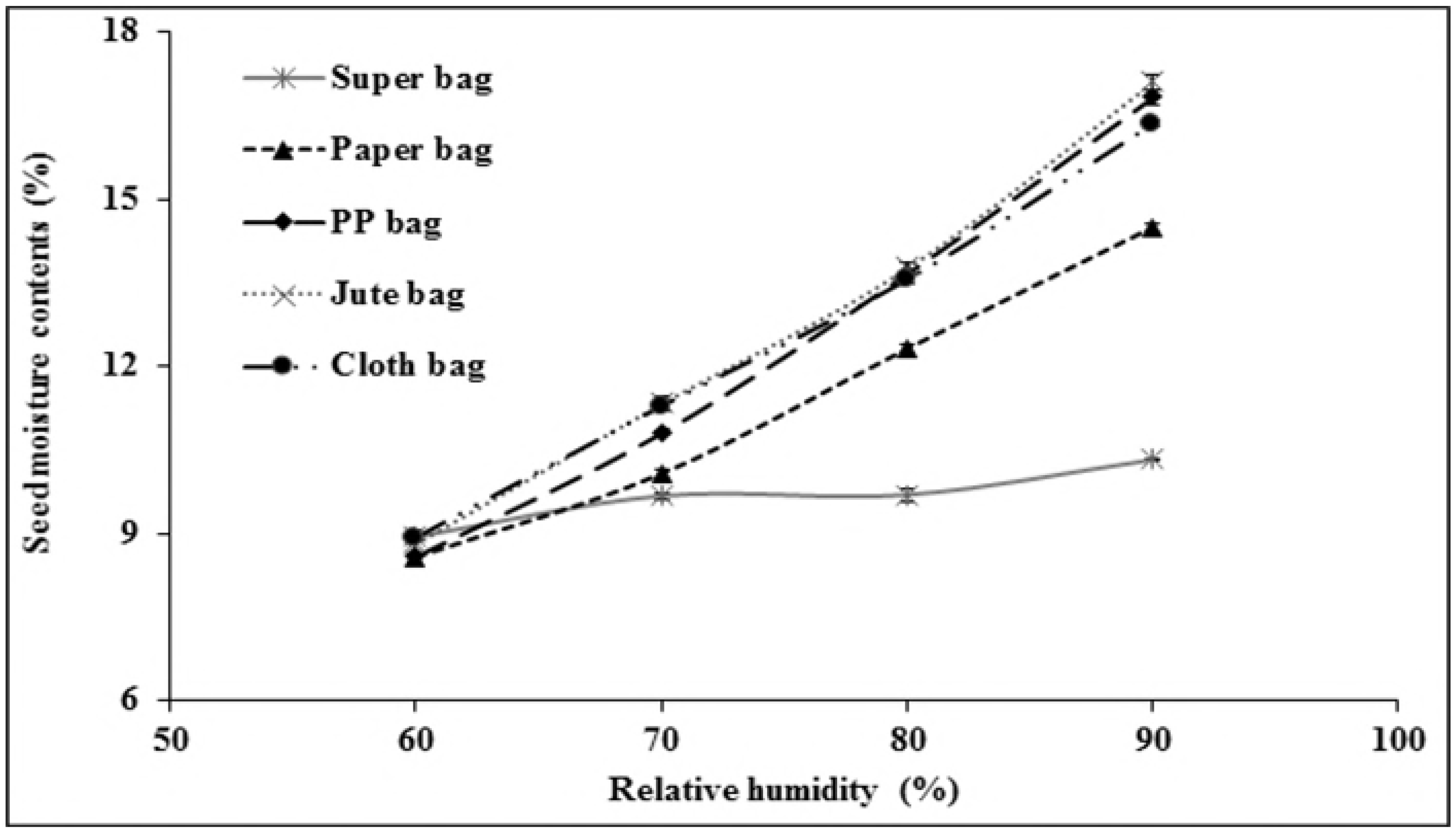
Seed moisture adsorption isotherms of wheat seeds in different packaging materials at 25°C. At 60% RH, maximum cotton seed moisture was recorded in cloth bag while minimum seed moisture was observed in PP and paper bags. Seeds in Super Bag had minimum moisture contents compared the seeds in all other packaging materials at 70% RH (Fig 4). Moisture contents of cotton seeds at 80% RH were significantly higher in cloth bag while Super Bag had seeds with lowest moisture. Cotton seed moisture also increased in Super Bag at 90% RH but still considerably low (10.39%) as compared to seed moisture in rest of packaging materials (Fig 4). Seed in PP bag had maximum seed moisture (17.52%) that was statistically similar to cloth bag (17.26%) and jute bag (17.20%) followed by paper bag (15.43%).

**Fig 4.**
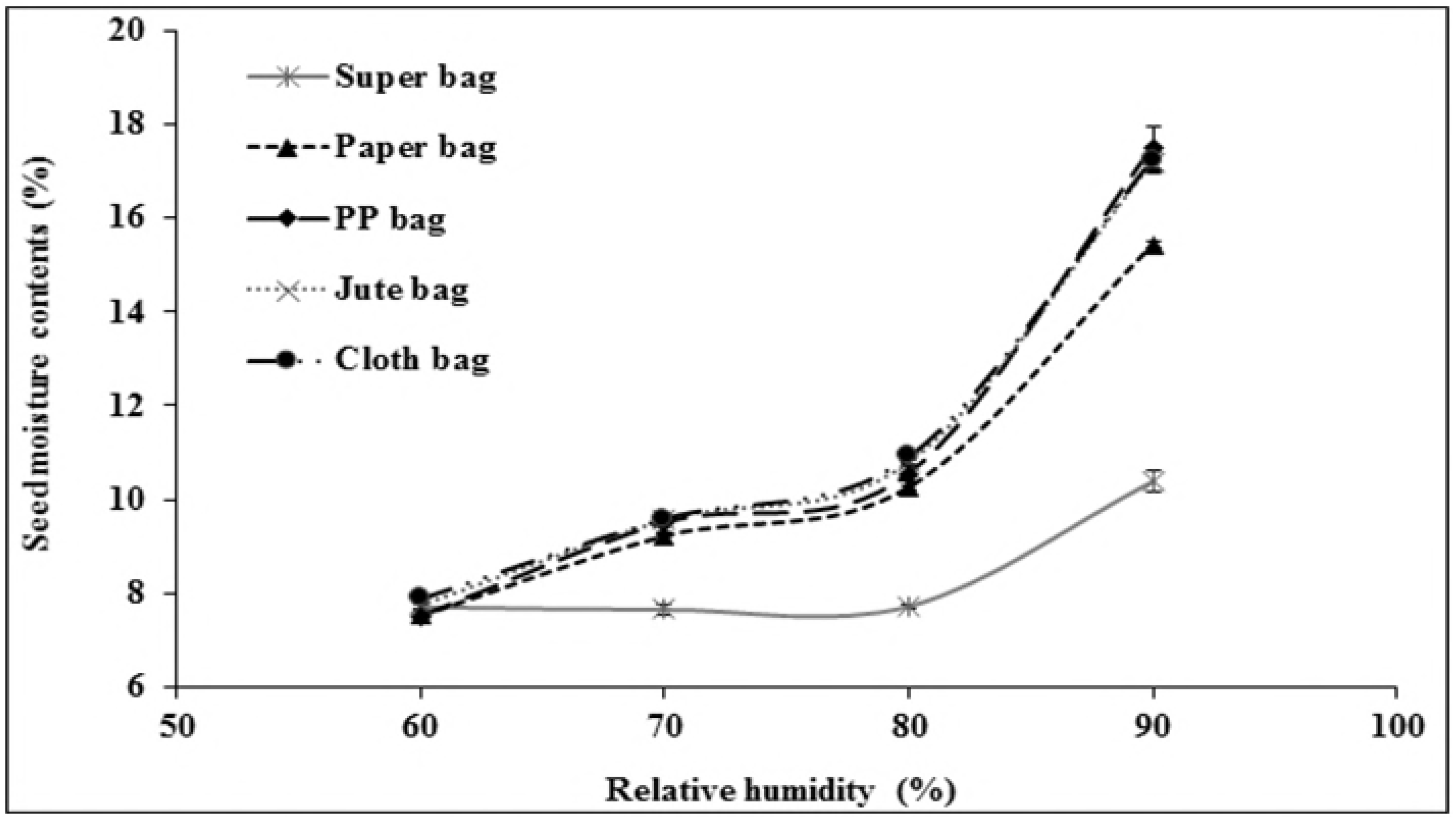
SeEd moisture adsorption isotherms of cotton seEds in different packaging materials at 25°C. There was a considerable difference in moisture contents of quinoa seeds placed in different packaging material at 60% RH. Seeds in Super Bag had minimum moisture contents compared to the seeds in all other storage materials (Fig 5). At 70% RH, Super Bag had seeds with minimum moisture contents while seed in paper, jute, cloth and PP bag had high moisture. Moisture contents of quinoa seeds at 80% RH were in order of Super Bag < paper bag < jute bag < cloth bag < PP bag (Fig 5). At 90% RH, seed moisture contents increased from 8.30% (at 80% RH) to 9.81% in Super Bag but compared to other bags this increment was far less than that was noted in other bags. Seed in jute bag had maximum seed moisture (17.25%) followed by PP bag (17%) and cloth bag (16.73%).

**Fig 5.**
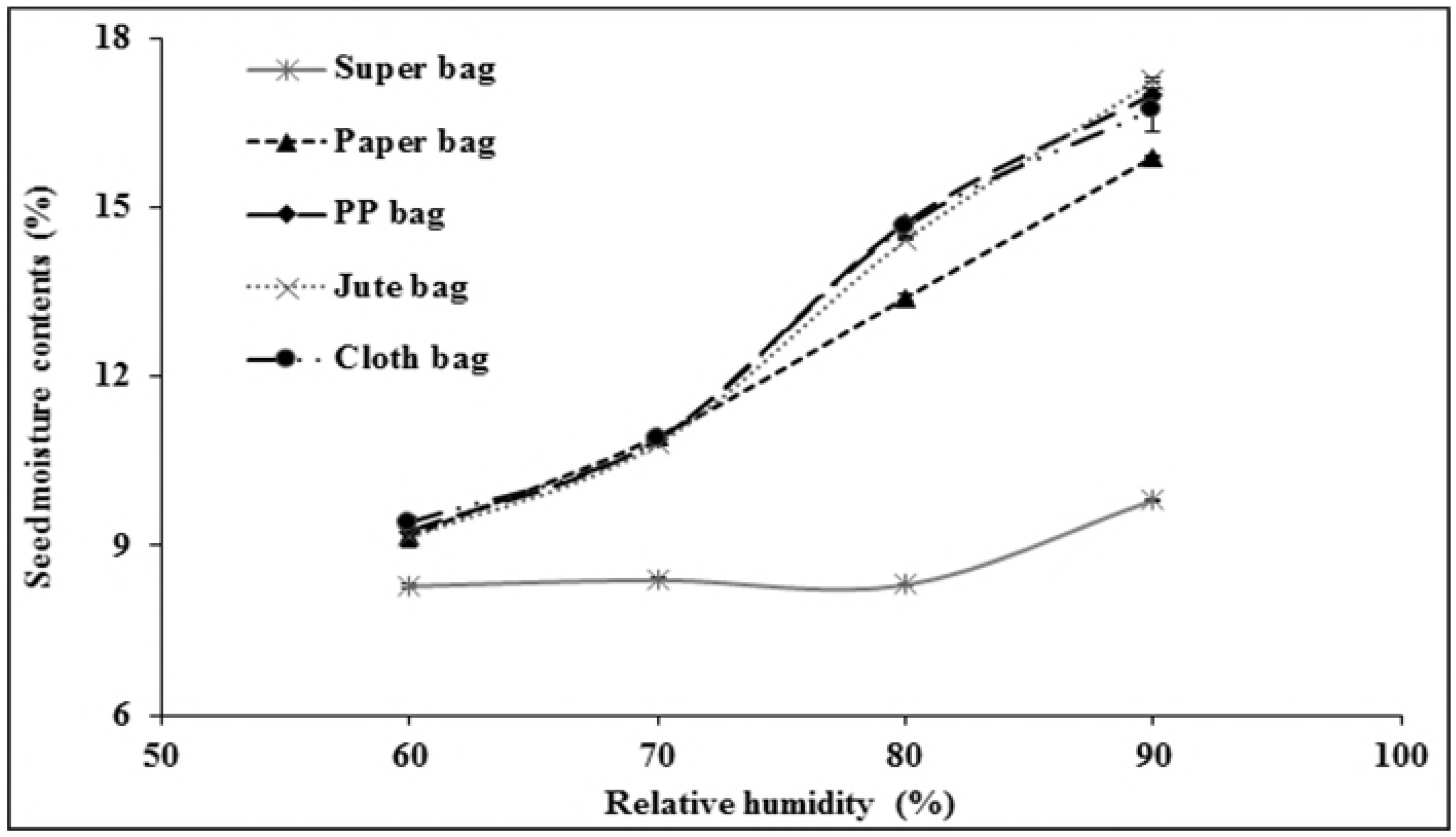
Seed moisture adsorption isotherms of quinoa seeds in different packaging materials at 25°C.

Seed deterioration and aging is a continuous process and its ultimate effects are loss of seed viability and vigor. High seed moisture content is the major culprit that can speed up the process of seed deterioration. High germination in Super Bag at 8% initial SMC was the outcome of Dry Chain that was continuously maintained by the hermetic nature of Super Bag. Reduction of seed moisture by initial drying and packaging in moisture proof containers can reduce the deterioration of commodities by fungi, storage losses due to insects and thus increase the shelf life seeds [1, 6]. Seeds stored in Super Bag at 14% initial SMC rapidly lost their germination as they were unable to lose their moisture due to barrier layer of the Super Bag and those continuous high moisture contents proved lethal for the seed germination. Major reason for the reduction of seed germination in conventional packaging materials could be the high seed moisture contents that lead to the production of reactive oxygen species causing oxidative damage and seed deterioration [20]. Fluctuations in RH of the storage environment affect seed moisture contents as is evident from our seed moisture adsorption isotherm study. Seed absorb moisture under conditions of high ambient relative humidity and quickly lost its germination due to the deteriorative process occurring at faster rate [21].

## Conclusion

Hermetic Super Bag maintained seed moisture up to 70% RH. At higher level of RH, moisture contents increased to some extent (1-2%) in Super Bag, whereas this increase was much higher in conventional packaging materials (≈9% higher than original SMC at 90% RH). Seed storage in Super Bag can help to prevent the significant increase in seed moisture at higher RH that will ultimately help to maintain high seed quality by implementing The Dry Chain.

## Acknowledgements

Authors are highly thankful to Higher Education Commission Pakistan, Pioneer Pakistan Seed, Wheat Research Institute AARI Pakistan, Rhino Research Group, Thailand and GrainPro, Inc. USA for support in provision of research materials and analysis facilities.

